# A Single-cell Spatiotemporal Atlas Encoding Age-dependent Evolution of the Ischemic Penumbra

**DOI:** 10.64898/2026.09.08.750251

**Authors:** Chenghao Jin, Li Fan, Qing Ye, Hongjian Pu, Wenting Zhang, Guanchen He, Junxuan Lyu, Wan-Chen Wu, Xiaoming Hu, Rehana K. Leak, Ke-Jie Yin, Kong Chen, Yejie Shi, Jun Chen

## Abstract

Aging is the predominant risk factor for stroke and a major driver of poor neurological recovery. Although the ischemic penumbra represents a key therapeutic target, its age-dependent molecular landscape remains poorly understood. Here, we generated a spatiotemporal transcriptomic atlas of the post-stroke penumbra in young (3-4 months) and aged (19-20 months) mice at acute (day 3) and chronic (day 14) stages following transient focal cerebral ischemia. Integrating spatial transcriptomics with dissociated single-cell RNA sequencing revealed a persistent transcriptomic penumbra (TP) that remains detectable for at least 14 days after stroke in both age groups, far beyond the window suggested by conventional imaging modalities. Aging profoundly altered TP organization, resulting in dysregulated energy metabolism, impaired neurovascular repair, diminished pro-reparative cell-cell communication, and extensive remodeling of transcriptional regulatory networks. Using this atlas, we further identified regulatory hub genes whose predicted perturbation could simultaneously counteract multiple age-associated pathological pathways. These data define the molecular architecture of the aging post-stroke penumbra and provide a publicly accessible framework for mechanistic discovery and therapeutic target prioritization aimed at improving recovery after stroke.

## Introduction

Ischemic stroke is the leading cause of neurological mortality and disability, imposing an enormous global public health burden worldwide^1,2^. Age is the strongest irreversible independent risk factor for stroke. Epidemiological data show that over 75% of ischemic strokes occur in people aged 65 and older^3,4^. Elderly patients exhibit larger infarcts, worse neurological injury, compromised repair capacity, and poorer clinical outcomes than young individuals^5,6^. The penumbra is the hypoperfused tissue around the infarct core. While it has electrophysiological dysfunction from reduced blood flow, it can be salvaged if the ion pump function is preserved until reperfusion and injured tissue can be repaired. These factors mitigate the secondary injury caused by oxidative DNA damage, excitotoxicity, spreading depression, and inflammation^7^. Hence, the penumbra is a major target for ischemic stroke therapies, determining the efficacy of clinical thrombolysis^8^/thrombectomy^9–12^ and shaping long-term neurological prognosis^13,14^. The stroke penumbra was initially defined based on the degree of loss of cerebral blood flow^15,16^ and subsequently by cellular phenotypic delineation based on neurovascular unit (NVU) structure^17^ and molecular definitions involving markers like heat shock protein 70 (HSP70)^18^. However, current criteria fail to capture the transcriptomic and cellular molecular heterogeneity inherent within the ischemic penumbra. Consequently, the key molecular signatures of the penumbra remain uncharacterized, and the impact of aging on its spatiotemporal dynamics remains unknown, creating an impediment to our understanding of the pathophysiological processes that drive poor outcomes in elderly stroke patients.

Spatial transcriptomics enables in situ, high-precision mapping of molecular and cellular architecture in fresh-frozen or formalin-fixed and paraffin-embedded tissues such as the brain^19–22^, but the stroke field still lacks a comprehensive spatial transcriptomic resource spanning young/aged and acute/chronic disease stages. Current studies are limited to young animals, single time points, or dissociated (non-spatial) single-cell RNA sequencing (scRNA-seq)^23–27^. Thus, the available literature does not discriminate penumbra-specific gene expression and cell-cell communication patterns, nor does it systematically characterize the perturbations induced by aging and disease progression. While the ischemic penumbra is defined through hemodynamic^15^, cellular^17^, and molecular^18^ criteria, no study has established a transcriptomic penumbra (TP) or dissected how aging and disease progression disrupt penumbral energy metabolism, synaptic function, endogenous repair functions, and immune homeostasis at the spatially discrete molecular level. Thus, the absence of a standardized spatial atlas of the stroke transcriptome as a function of age as well as the passage of time (injury stage) severely restricts mechanistic research and the development of targeted therapies for ischemic stroke.

Here, we present the first cross-age, cross-stage spatiotemporal transcriptomic resource for ischemic stroke using 10X Visium. We evaluated 20 mouse brains from young and aged mice at sham, acute (day 3), and chronic (day 14) post-stroke stages, leveraging 62,958 high-quality Visium spots. We define and comprehensively characterize the TP and decipher its modular dynamics during ischemic injury progression and aging-dependent perturbations. By integrating spatial transcriptomic and scRNA-seq data, we identify aging-sensitive cell types, impaired reparative ligand-receptor interactions, and core transcription factors driving age-related dysfunction in the penumbra. This new dataset thus represents the first publicly available spatiotemporal atlas of ischemic stroke as a function of age, providing a high-quality resource for dissecting mechanisms that exacerbate stroke and advancing precision therapy for young as well as elderly patients.

## Results

### Landscape of the dynamic stroke transcriptome across spatial and temporal dimensions

We collected ipsilateral coronal brain slices from young (3–4 month) and aged (19–20 month) male mice, randomized into sham-operation or transient middle cerebral artery occlusion (tMCAO) groups and euthanized at acute (3 days) or chronic (14 days) stages (**Fig. 1a**). Post-stroke magnetic resonance imaging (MRI) verified successful induction of the ischemic stroke model (**Supplementary Fig 1**). Using the 10x Visium^28^ spatial transcriptomic platform, we profiled 62,958 high-quality spots from 20 mice with an *n* of 3-4 per group (**Fig. 1a–c**, **Supplementary Table 1**).

**Figure 1.**
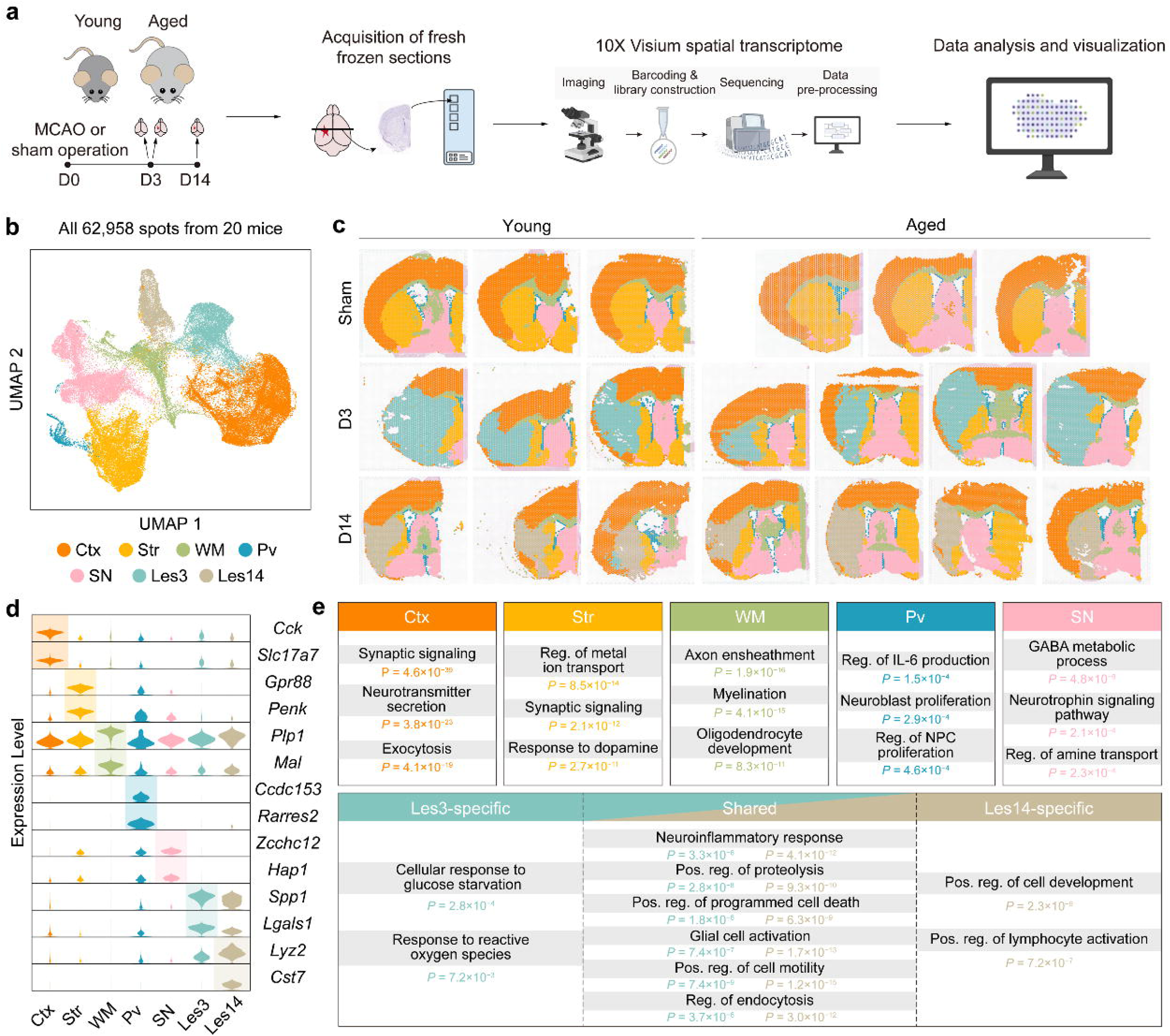
Establishment of a spatiotemporal transcriptomic atlas in ischemic stroke brains of young and aged mice. **(a)** Schematic workflow of sample preparation, 10x Visium spatial transcriptomic detection, and bioinformatic analysis. Young and aged mice were subjected to the sham operation or tMCAO surgery. Fresh-frozen brain tissues were collected from sham-surgery controls (n=3 for both young and aged), or D3 (n=3 for young and n=4 for aged), and D14 post-stroke (n=3 for young and n=4 for aged) for sequencing. **(b)** UMAP visualization of 62,958 high-quality spatial spots from 20 mice. Spots are color-labeled by anatomical and lesion clusters, including Ctx, Str, WM, Pv, SN, Les3, and Les14. **(c)** In situ spatial distribution of all annotated clusters in (b) across young and aged groups at sham, D3, and D14 stages. **(d)** Violin plots displaying expression levels of representative marker genes for each cluster. **(e)** Representative GO BP enrichments for Ctx, Str, WM, Pv, and SN anatomical brain regions. Les3-specific, Les14-specific, and shared enriched GO BP terms between the two lesion clusters are presented. *P*-values for GO BP enrichment were calculated by Fisher’s exact test. Ctx: cerebral cortex; Sham: sham-surgery controls; D3: day 3 post-operation; D14: day 14 post-operation; GO BP: Gene Ontology biological process; Les3: lesion subregion 3; Les14: lesion subregion 14; tMCAO: transient middle cerebral artery occlusion; Pv: periventricular region; SN: septal nuclei; Str: striatum; UMAP: uniform manifold approximation and projection; WM: white matter.

Unsupervised clustering classified all spots into 7 distinct clusters (**Fig. 1b**). Based on spatiotemporal distribution (**Fig. 1c**) and cluster-specific marker profiles (**Fig. 1d**, **Supplementary Table 2**), we annotated these clusters as cortex (Ctx), striatum (Str), white matter (WM), periventricular area (Pv), septal nuclei (SN), day-3-specific lesion (Les3), and day-14-specific lesion (Les14). Spatial mapping across time points showed well-defined, anatomically consistent compartmentalization in young and aged mice, alongside robust stage-specific lesional signatures from acute to chronic stage (**Fig. 1c**). These regionally conserved spatial architectures, combined with the clustering patterns (**Fig. 1b**) and cluster-specific marker expression (**Fig. 1d**, **Supplementary Fig 2a**), verified robust separation of these 7 clusters.

To explore the biological functions of each cluster, we performed Gene Ontology (GO) enrichment analysis on the highly expressed genes (HEGs, **Fig. 1e**, **Supplementary Table 2**). Consistent with known forebrain physiology, Ctx and Str were enriched in synaptic signaling and neurotransmitter secretion; WM in axon ensheathment and myelination; Pv in neuroblast proliferation and IL-6 production; and SN in GABA metabolism and neurotrophin signaling (**Fig. 1e**). These region-specific patterns verify the authenticity of this spatial transcriptomic dataset.

For lesional clusters Les3 and Les14, we observed pervasive ribosomal gene upregulation and excluded it from functional analysis to avoid signal masking (**Supplementary Fig 2b–e**). GO analysis of non-ribosomal HEGs showed the shared biological processes (BPs) covering neuroinflammation, glial activation, proteolysis, programmed cell death, cell motility, and endocytosis, suggesting a spectrum of activated functions in the lesional area across ischemic stages (**Fig. 1e**). Les3 was uniquely enriched in glucose starvation and reactive oxygen species, while Les14 was characterized by cell development and lymphocyte activation (**Fig. 1e**), suggesting acute cellular response activation at 3 days, followed by chronic endogenous repair/adaptive immune remodeling at 14 days.

In summary, we established a high-resolution spatiotemporal transcriptomic landscape of ischemic stroke across age and injury stage. This dataset defines canonical anatomical compartments and captures dynamic, stage-specific lesional signatures, thus establishing a reliable resource for exploration of stroke pathophysiology in the context of aging, the passage of time post-injury, neural cell type, anatomical subregion, and cell-cell communication patterns.

### Defining the transcriptomic penumbra (TP) through specific molecular signatures

Preliminary clustering identified lesion clusters (Les3, Les14) but failed to capture their internal heterogeneity. We thus performed secondary clustering on Les3 and Les14 spots, revealing a consistent ‘periphery-around-core’ spatial architecture across young and aged ischemic mice at both the acute and chronic stages: Les3 were divided into 5 subclusters (2 peripheral [Peri], 3 core [Core]), while Les14 comprised 4 subclusters (2 Peri, 2 Core) (**Fig. 2a–d**, **Supplementary Fig. 3a–d**). This ‘periphery-around-core’ structure was present in all post-stroke samples (**Fig. 2c–d**, **Supplementary Fig. 3c–d**).

**Figure 2.**
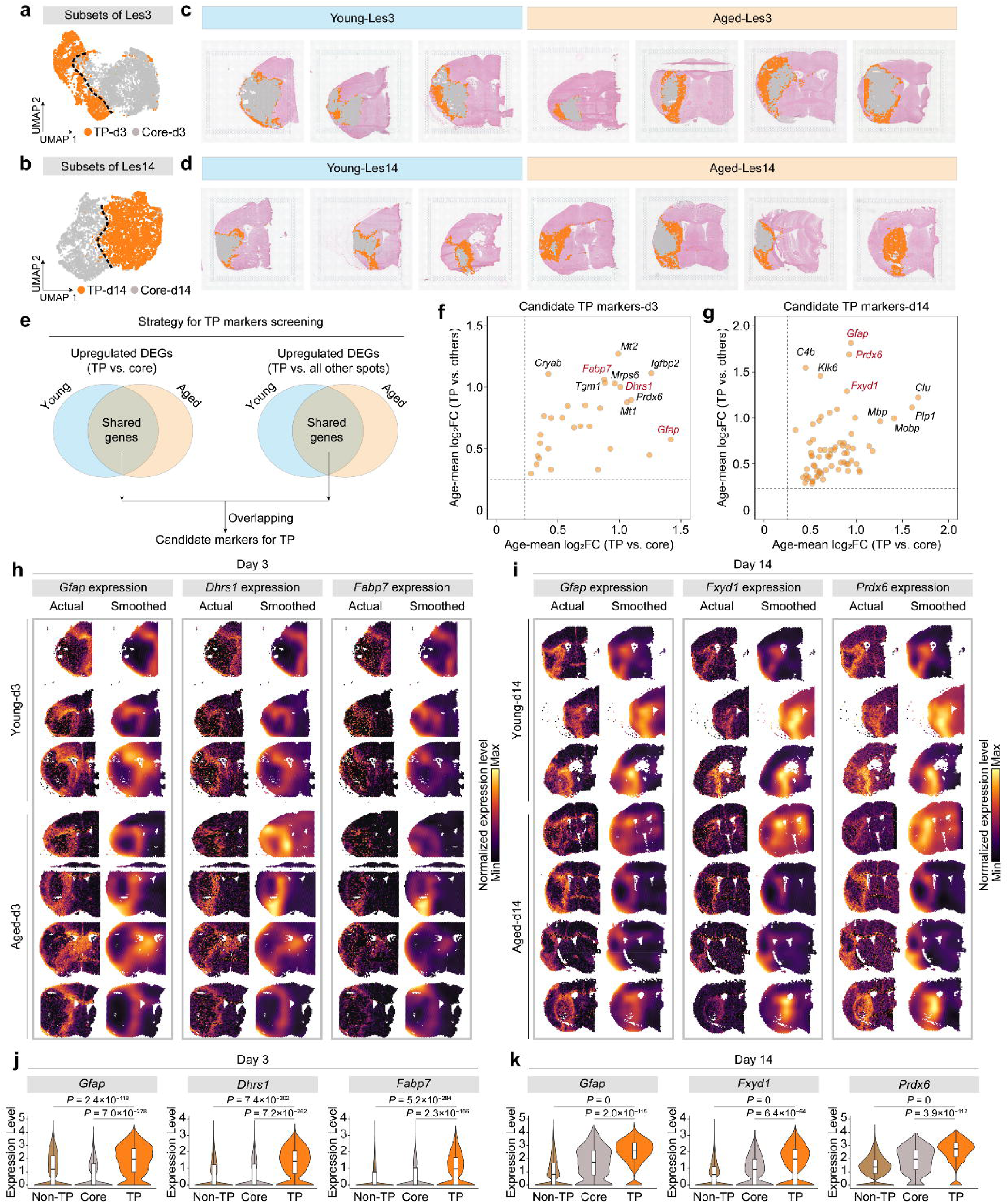
Identification and spatiotemporal profiling of transcriptomic penumbra-specific markers in ischemic stroke. **(a–b)** Subsets of Les3 and Les14 clusters, containing TP and Core subregions at acute D3 stage (a) and chronic D14 stage (b). **(c–d)** In situ spatial distribution of TP and Core spots at D3 (c) and D14 (d) in young and aged post-tMCAO mice. **(e)** Schematic strategy for TP-specific marker screening. **(f–g)** Candidate TP marker genes at acute D3 stage (f) and chronic D14 stage (g). **(h–i)** Visualization of actual and smoothed expression levels of representative TP markers at D3 (h) and D14 (i) across all samples. **(j– k)** Violin plots comparing marker expression levels among non-TP, Core, and TP regions at D3 (j) and D14 (k). *P*-values were calculated by Wilcoxon rank-sum tests with Bonferroni correction. D3: day 3 post-operation; D14: day 14 post-operation; Les3: lesion subregion 3; Les14: lesion subregion 14; tMCAO: transient middle cerebral artery occlusion; TP: transcriptomic penumbra.

Spatially, the Peri region occupied an intermediate anatomical position between the Core and non-lesional brain tissues (**Fig. 2a–d**, **Supplementary Fig. 3a–d**). Registration with T2-weighted MRI and H&E histological staining further confirmed that the Peri region localized to the peri-infarct zone, defined as radially extending 200 to 300 microns from the infarct core (**Supplementary Fig. 3e**). Transcriptionally, Uniform Manifold Approximation and Projection (UMAP) visualizations showed distinct topological boundaries between Core and Peri spots in both Les3 and Les14 clusters (**Fig. 2a–b**, **Supplementary Fig. 3a–b**). Approximately 1,000 differentially expressed genes (DEGs) were identified between Core and Peri in each post-stroke group (**Supplementary Fig. 3f**). Consistent with the classical definition of the ischemic penumbra established by imaging and perfusion studies^15,16^, we designated this Peri region as the Transcriptomic Penumbra (TP).

The salvageable penumbra is the primary therapeutic target for stroke intervention, but its molecular characteristics remain poorly defined. Using a conservative screening pipeline (**Fig. 2e**), we overlapped age-insensitive upregulated DEGs from two sets, including TP-specific genes against core lesions and non-TP areas. We obtained 30 acute-stage and 70 chronic-stage TP candidate markers (**Fig. 2f–g**, **Supplementary Table 3**). Notably, only 6 genes (*Cryab, Gfap, Metrn, Mt1, Prdx6, S100a16*) were shared between stages (**Supplementary Table 3**), supporting our determination of stage-specific TP signatures and revealing the dynamic evolution of the TP with the passage of time after ischemic injury.

We further validated TP candidate markers by assessing their spatial distribution (**Supplementary Fig. 4**) and prioritized optimal markers. *Gfap* encodes glial fibrillary acidic protein (GFAP), a classic astroglial scar marker^29^, which was specifically expressed in TP at both 3 and 14 days post-stroke (**Fig. 2h–k**). We also identified novel stage-specific TP markers. *Dhrs1* and *Fabp7* showed acute-stage-specific expression (**Fig. 2h**), while *Fxyd1* and *Prdx6* were specific to the chronic injury stage (**Fig. 2i**). Statistical analysis confirmed their significantly higher expression in TP than Core and non-TP areas (**Fig. 2j–k**).

Functionally, fatty acid-binding protein 7 (FABP7) has been reported to be markedly upregulated in the ischemic penumbra at 6–48 h after stroke^30,31^, aligning with our observations at 3 days. It worsens neurological damage by regulating mitochondrial function in the acute stage^30,31^ but may also promote neural regeneration and repair at the subacute stage (7 days post-stroke)^32^. Peroxiredoxin 6 (PRDX6) is highly expressed in astrocytes after ischemia; its calcium-independent phospholipase A2 (iPLA2) activity promotes neuroinflammation and neuronal apoptosis^33,34^, consistent with findings that extracellular peroxiredoxin family proteins act as damage-associated molecular patterns (DAMPs) to drive post-ischemic inflammation via TLR2/4 signaling^35,36^. In addition, Prdx6 also alleviates oxidative stress injury post-stroke^37^, supporting its potential dual regulatory effects following ischemia^38^. Dehydrogenase/reductase SDR family member 1 (DHRS1) is involved in regulating mitochondrial energy metabolism and oxidative stress^39^. As a regulatory subunit of Na /K -ATPase, FXYD domain containing ion transport regulator 1 (FXYD1) modulates neuronal excitability and potassium homeostasis^40^. Although both are widely expressed in the brain^41,42^, their post-stroke roles remain largely uncharacterized.

We further validated the spatial specificity of these candidate TP markers at the protein level using immunofluorescence staining (**Supplementary Fig. 5**). Four markers (GFAP at both day 3 and day 14, FABP7 at day 3, PRDX6 and FXYD1 at day 14) displayed well-restricted spatial localization within the TP. In contrast, despite specific upregulation of *Dhrs1* mRNA in the acute TP, its protein distribution failed to recapitulate this spatial specificity, likely reflecting a temporal decoupling between mRNA transcription and protein translation.

In summary, we defined TP and characterized its stage-specific molecular signatures in ischemic lesions, advancing the molecular understanding of the ischemic penumbra.

### Modular dissection of temporal and functional dynamics of the TP of young ischemic mice

To dissect temporal molecular dynamics and functional alterations in the TP, we performed pairwise transcriptomic comparisons among the sham hemisphere, TP at 3 days, and TP at 14 days in young mice. A total of 2,631 time-varying genes (TVGs) were identified (**Supplementary Table 4**) and classified into 16 distinct expression modules based on their temporal patterns across groups (**Fig. 3a–b**, **Supplementary Table 5**). The average expression patterns of each module were validated to match the predicted temporal dynamics (**Fig. 3c**). We further grouped these 16 modules into 5 functional categories based on consistent expression trajectories. All 16 modules were further grouped into five functional clusters according to expression characteristics: suppressed modules (9, 10, 14, 12), activated modules (2, 1, 13, 11), bounced-back modules (7, 8), fallen-back modules (4, 3), and over-corrected modules (6, 15, 5, 1. 16) (**Fig. 3c**).

**Figure 3.**
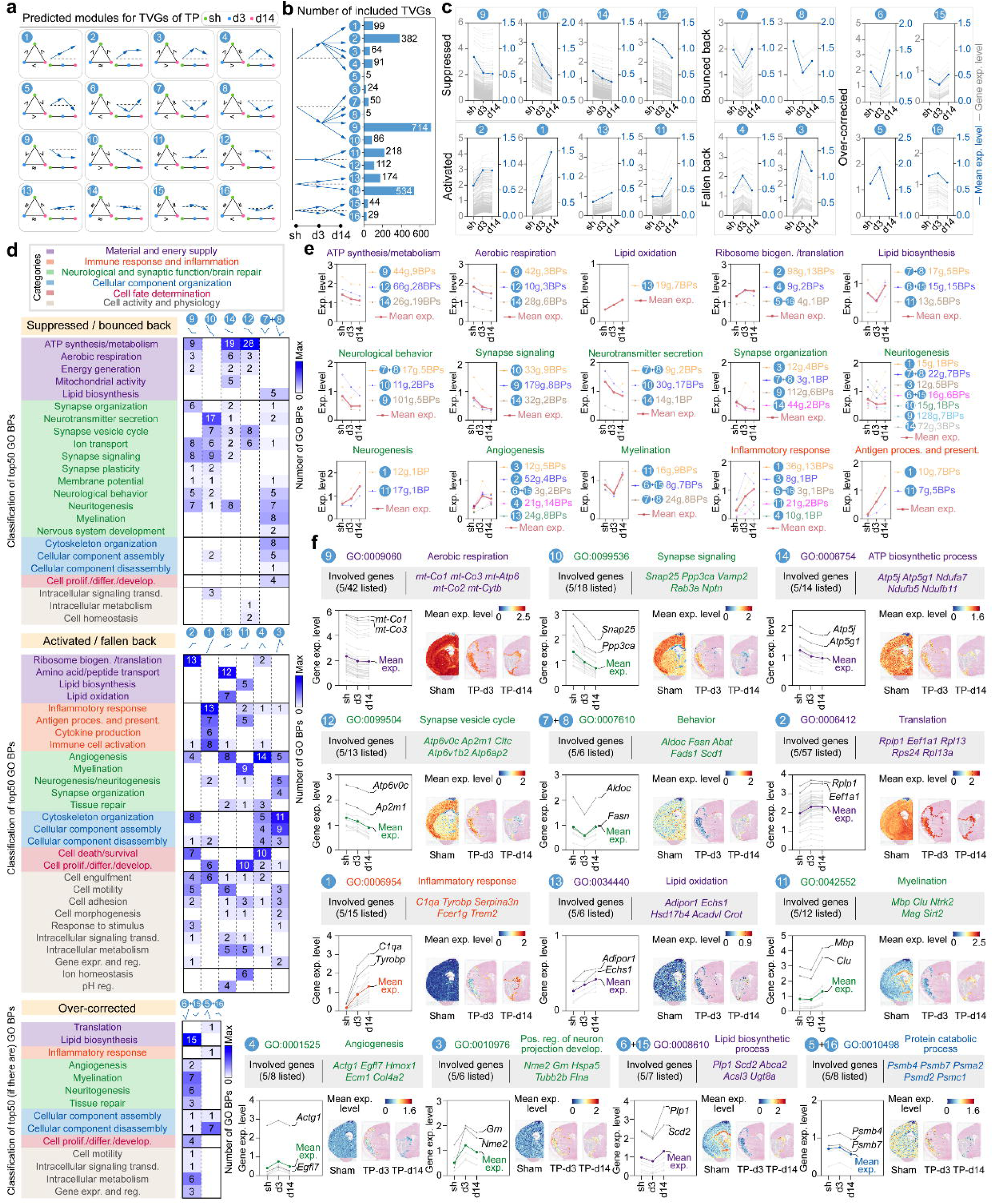
Spatiotemporal dynamic module identification and functional profiling of time-varying genes in TP of ischemic stroke. **(a)** Schematic diagrams illustrating dynamic expression relationships among sham, D3, and D14 groups for TVGs in each predicted module. **(b)** Bar plot displaying the number of TVGs in each of the 16 modules. **(c)** Individual temporal expression trajectories of all involved TVGs (grey lines) and module-averaged expression trends (blue lines) across modules. The 16 modules are categorized into 5 dynamic types: suppressed, activated, bounced back, fallen back, and over-corrected. **(d)** Heatmap showing classification of top 50 significant GO BPs enriched from TVGs of each module, grouped into 6 major categories and ∼40 minor categories. **(e)** Modular expression levels of 15 key minor GO BP categories, with bold red lines representing the average expression levels. **(f)** The top 5 highly expressed genes involved in each representative GO BP for each module. Grey lines show expression of all involved genes; colored lines show mean expression levels. Spatial visualization of mean expression levels for all involved genes is additionally presented. *P*-values for GO BP enrichment were calculated by Fisher’s exact test. D3: day 3 post-operation; D14: day 14 post-operation; GO BP: Gene Ontology biological process; tMCAO: transient middle cerebral artery occlusion; TP: transcriptomic penumbra; TVGs: time-varying genes.

GO enrichment analysis on the TVGs was conducted to decode module-related functions. Among them, limited gene sizes restricted reliable enrichment for Modules 5, 6, and 8; thus, trajectory-consistent modules with identical trajectory categories were merged for subsequent analysis. These sets include Module 5 plus 16, Module 6 plus 15, and Module 7 plus 8. We next conducted GO BP enrichment analysis for all individual and combined modules, and classified the top 50 significant GO terms from each module into 6 major functional categories, including material and energy supply, immune response and inflammation, neurological and synaptic function and brain repair, cellular component organization, cell fate determination, and cell activity and physiology. Around forty minor categories were further classified based on functional similarities and the genes involved (**Fig. 3d**, **Supplementary Table 6**), revealing the complex dynamic reprogramming of TP biology after stroke. Given that GO terms associated with each minor functional category were distributed across multiple temporal modules (**Fig. 3d**), their collective dynamics could only be fully captured by integrating across these modules. We thus quantified the integrated temporal trends of 15 key minor functional categories across modules (**Fig. 3e**). We then selected 13 representative GO BP terms to dissect functional dynamics at the gene level (**Fig. 3f**). Temporal expression patterns of involved genes within each selected GO term were consistent with their respective module trajectories, and spatial projection analysis visualized functional shifts in TP, with annotated signature genes of each process (**Fig. 3f**).

As expected, based on prior definitions of the stroke penumbra^7^, respiration and energy metabolism pathways showed prominent suppression, including ATP synthesis/metabolism and aerobic respiration, with continuous downregulation across stroke progression (**Fig. 3d–e**). The most prominent genetic drivers of the observed downregulation involved mitochondrial genes (*mt-Co1/2/3*, *mt-Atp6*, etc.) and ATPase subunit genes (*Atp5j*, *Atp5g1*, etc.) and respiratory complex I subunits (*Ndufa7*, *Ndufb5*, etc.) (**Fig. 3f**). In contrast, lipid oxidation processes were progressively upregulated in activated Module 13, driven by *Adipor1* (encoding adiponectin receptor 1, a crucial regulator of lipid oxidation^43^) and several genes encoding fatty acid β-oxidation enzymes (*Echs1*, *Hsd17b4*, etc.) (**Fig. 3d–f**). These results indicate impaired energy supply in TP after stroke, with compensatory lipid metabolic reprogramming as an alternative energy source.

Neurological and synaptic function processes were predominantly enriched in the suppressed modules, including synapse organization, neurotransmitter secretion, neurological behavior, and neuritogenesis (**Fig. 3d**). Synaptic function remained impaired at day 14 after stroke. Multiple core synaptic regulatory genes (*Ap2m1*, *Atp6v0c*, etc.) and vesicle-associated genes (*Snap25*, *Vamp2*, etc.) drove these suppressions (**Fig. 3e**). Given that intact synaptic function is essential for adaptive behaviors, these suppressed synapse-related BPs may contribute to impaired neurological function after stroke. However, partial synaptic rebound processes were enriched in bounced-back modules, and the corresponding enzyme-encoding genes (*Aldoc*, *Fasn*, etc.) mediated this partial functional recovery after stroke (**Fig. 3d–f**).

Ribosome biogenesis and translation processes remained upregulated after stroke, driven by abundant ribosomal subunits (*Rplp1/13/13a*, *Rps24*, etc.) and translation elongation gene (*Eef1a1*), reflecting elevated demand for protein synthesis in TP (**Fig. 3d–f**). Correspondingly, protein catabolic processes were suppressed in the chronic phase (Module 5 plus 16), with marked reductions in the expression of proteasome-related genes (*Psma2/b4/b7/c1/d2*). These findings suggest homeostatic shifts from protein degradation to protein synthesis in the TP after stroke.

Endogenous repair processes were broadly activated in TP across multiple modules. Neurogenesis and neuritogenesis processes were temporally upregulated following stroke, with genes encoding regulators of neuronal development and growth (*Grn, Nme2*, etc.) being dominantly involved (**Fig. 3d–f**). Angiogenesis-related processes, including endothelial cell (EC) proliferation, EC migration, and tube formation (**Supplementary Table 6**), reflect the precise, programmed regulation of angiogenesis (**Fig. 3d**). Especially in the acute stage, genes encoding angiogenic regulators (*Actg1*, *Egfl7*, etc.) were highly expressed (**Fig. 3e–f**). Myelination and lipid biosynthesis pathways showed acute suppression followed by chronic activation, with dominant genes involved encoding key myelin components (*Mbp*, *Mag*, etc.), myelination modulators (*Sirt2*, *Ntrk2*, etc.), and myelin/lipid synthesis (*Plp1*, *Scd2*, etc.) (**Fig. 3d–f**), aligning with lipid composition characteristics of myelin sheaths^44^.

Immune and inflammatory responses were mainly gathered in activated Module 1 (**Fig. 3d**), showing continuous progressive upregulation after stroke, including immunoinflammatory response and antigen processing and presentation (**Fig. 3e**). Core driving genes include major neuroinflammatory mediators (*C1qa*, *Tyrobp*, etc.) (**Fig. 3f**), indicating progressive immune activation and sustained inflammatory stress in TP after stroke.

In summary, this comprehensive modular analysis unveils temporal reprogramming in young mouse TP after ischemic stroke. Mechanistically, the TP exhibits energy deficiency and synaptic impairment, coupled with coordinated activation of lipid metabolism, protein synthesis, endogenous repair cascades, and chronic neuroinflammation. Systematic screening of function-specific signature genes also provides valuable molecular candidates for targeted modulation of TP pathophysiology.

### Modular dissection of aging-driven spatiotemporal perturbations in the TP

To elucidate aging-induced TP perturbations, we first compared the differences in size of TP between the young and aged groups (**Supplementary Fig. 6a–b**). Surprisingly, neither the proportion of TP area nor the density of TP spots in the ipsilesional hemisphere were smaller as expected (see Discussion) in the aged compared to young group. Rather, the aged TP was larger in the chronic phase (**Supplementary Fig. 6b**), demonstrating that salvageable areas still exist in the post-stroke aged brain. Next, we systematically compared spatiotemporal transcriptomic profiles between young and aged TP. UMAP projections revealed obvious topological shifts of TP cell distribution between young and aged mice (**Fig. 4a**). We identified 295 and 229 age-associated DEGs at 3 days and 14 days in TP, respectively (**Fig. 4b**, **Supplementary Table 7**). Using the same modular classification pipeline applied to young TP, we generated 16 consistent expression modules for aged TVGs (**Supplementary Table 8–9**). Cross-age module comparison further identified age-sensitive TVGs (ASTVGs) (**Supplementary Table 10**), which were broadly distributed across all modules (**Fig. 4c**). These findings indicate that aging profoundly rewires temporal transcriptomic dynamics within the TP.

**Figure 4.**
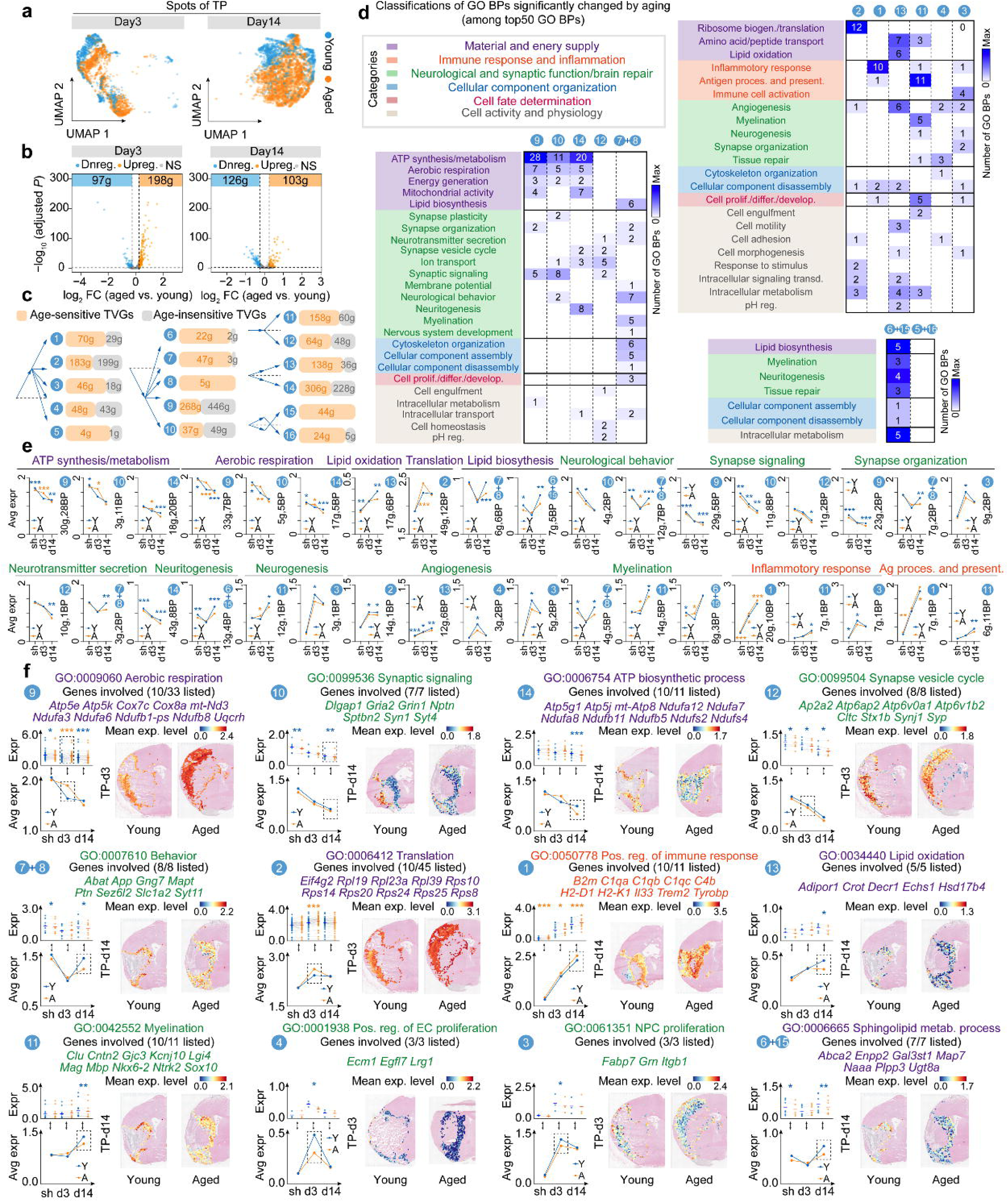
Age-dependent dynamic module and functional profiling of age-sensitive time-varying genes in TP of ischemic stroke. **(a)** UMAP visualization of TP spot distribution shifts between young and aged groups at D3 and D14 stages after stroke. **(b)** Volcano plots showing DEGs between young and aged TP at D3 acute and D14 chronic stages. **(c)** Numbers of ASTVGs and age-insensitive TVGs in each dynamic module. **(d)** Heatmap showing classification of top 50 significant GO BPs enriched from module-specific ASTVGs. **(e)** Comparisons of modular expression levels for representative GO BP categories between young (blue lines) and aged (yellow lines) TP. **(f)** The top representative involved genes of the GO BP term in each module. Scatter plots and line graphs show age differences in gene expression (up) and average expression levels (down). Spatial visualizations show age differences in mean expression levels of all involved genes. *P*-values for GO BP enrichment were calculated by Fisher’s exact test. *P*-values for DEG analysis were calculated by the paired t-test. ASTVGs: age-sensitive time-varying genes; D3: day 3 post-operation; D14: day 14 post-operation; DEGs: differentially expressed genes; GO BP: Gene Ontology biological process; tMCAO: transient middle cerebral artery occlusion; TP: transcriptomic penumbra; TVGs: time-varying genes; UMAP: uniform manifold approximation and projection.

Module-based GO enrichment analysis of ASTVGs revealed widespread functional disturbances associated with aging (**Fig. 4d, Supplementary Table 11**). Quantitative comparison of function-specific gene expression further demonstrated that aging exerts bidirectional effects on key biological processes (**Fig. 4e**). Aging predominantly suppressed pathways involved in energy and lipid homeostasis, neurological and synaptic function, and multiple endogenous repair programs, potentially contributing to impaired neurological recovery after stroke in older individuals. Notably, several GO pathways exhibited acute-phase upregulation in the aged TP, including ATP synthesis and metabolism, aerobic respiration, translation, behavior, neurogenesis, and myelination. Despite this apparent transcriptional activation, aged mice continued to exhibit marked functional deficits following stroke, suggesting that these responses may not be sufficient to restore tissue homeostasis or overcome age-related impairments. Aging also exerted bidirectional regulatory effects on immune pathways in the TP. Inflammatory responses and antigen-processing programs were enhanced in Module 1 of aged mice, whereas these pathways were suppressed in aged Module 11 (**Fig. 4e**). Collectively, these findings suggest that aging disrupts immune homeostasis and alters the coordination of adaptive and reparative responses in the TP.

We analyzed 12 representative aging-sensitive GO BPs to resolve functional perturbations at the gene level (**Fig. 4f**). Spatial expression patterns further visualized these age-specific functional alterations (**Fig. 4f**). We characterized the ASTVGs involved in these aging-sensitive functions. Compared to the young TP, aging impaired aerobic respiration and ATP synthesis in the chronic stage post-stroke by downregulating mitochondrial respiratory complex genes (Complex I subunits: *Ndufa3*, *Ndufa6*, etc.; complex III subunits: *Uqcrh*; cytochrome c oxidase subunits: *Cox7c*, *Cox8a*, etc.) and ATP synthase genes (*Atp5e*, *Atp5k*, etc.). Lipid oxidation pathways were suppressed by aging through reduced expression of fatty acid β-oxidation regulators (*Adipor1*, *Crot*, etc.). Aging promoted ribosome biogenesis and translation via upregulation of ribosomal subunit genes (*Rpl19/23a/39*, etc.). Neurological and synaptic functions were broadly suppressed with age through downregulation of core synaptic genes (*Syn1*, *Syt4/11*, etc.).

Aging broadly weakened endogenous repair programs (**Fig. 4f**), including neural precursor cell proliferation involving pro-neurogenesis genes (*Fabp7*, *Grn*, *Itgb1*), EC proliferation involving pro-angiogenesis genes (*Ecm1*, *Egfl7*, *Lrg1*), and myelination involving myelin-component genes (*Mag*, *Mbp*) and myelination regulatory genes (*Clu*, *Cntn2*, etc.), accompanied by suppressed sphingolipid metabolism (*Abca2*, *Enpp2*, etc.). Conversely, aging amplified the immune response through upregulation of MHC-I class molecule genes (*H2-D1*, *H2-K1*, *B2m*), complement molecule genes (*C1qa/b/c*, *C4b*), and immune inflammatory mediator genes (*Tyrobp*, *Trem2*, *Il33*). Notably, most ASTVGs overlapped with core driver genes identified in young TP (**Fig. 3f**), supporting the central nature of these regulatory genes, as they were highly vulnerable to aging-mediated dysregulation.

In summary, aging extensively reshapes the molecular dynamics of TP spatiotemporal profiles. We characterized aging-induced alterations in energy metabolism, synapse function, immune homeostasis, and endogenous repair processes. We have identified signature genes contributing to these age-dependent perturbations. These findings fill gaps in the mechanistic understanding of aggravated stroke outcomes in aged individuals.

### Integrated spatial and single-cell transcriptomic dissection of cellular aging landscapes in TP

To dissect TP cellular composition and age-induced transcriptomic alterations, we integrated our spatial transcriptomic data with our previously published scRNA-seq dataset from young and aged stroke mice^24^ (**Fig. 5a**). We performed robust cell type decomposition (RCTD)^45^ to infer cellular abundance weights within individual Visium spots. TP regions exhibited heterogeneous cellular compositions (**Fig. 5b–c, Supplementary Table 12**). Under sham conditions, five resident non-immune cell types—neurons (NEUR), oligodendrocytes (OL), choroid plexus epithelial cells (CPC), oligodendrocyte precursor cells (OPC), and astrocytes (ASC)—were the most abundant, collectively accounting for 92.9% of the total proportion, while immune cells were scarce, comprising only 2.1% of the total. Following stroke, these five resident non-immune cells remained the top six non-immune populations in TP. Notably, fibroblasts (FB) emerged as a newly dominant cell type in the TP, with their abundance significantly elevated from <1% to 12.9% at day 3 and 10.1% at day 14. Meanwhile, immune cells were markedly upregulated post-stroke, representing 15.4% at day 3 and 20.5% at day 14 of all cells. Among them, microglia and macrophages (MG/MΦ) were predominant cell types of the total immune cell pool, accounting for 84.0% at day 3 and 86.7% at day 14. Notably, RCTD-derived cellular weights, which reflect the proportion of the transcriptome of specific cells within the total transcriptome of a spot, may differ from actual cell counts due to variable transcript richness and cell volumes across CNS cell types (e.g., reported 5∼12% microglial proportion in CNS^46^ vs. lower RCTD weights).

**Figure 5.**
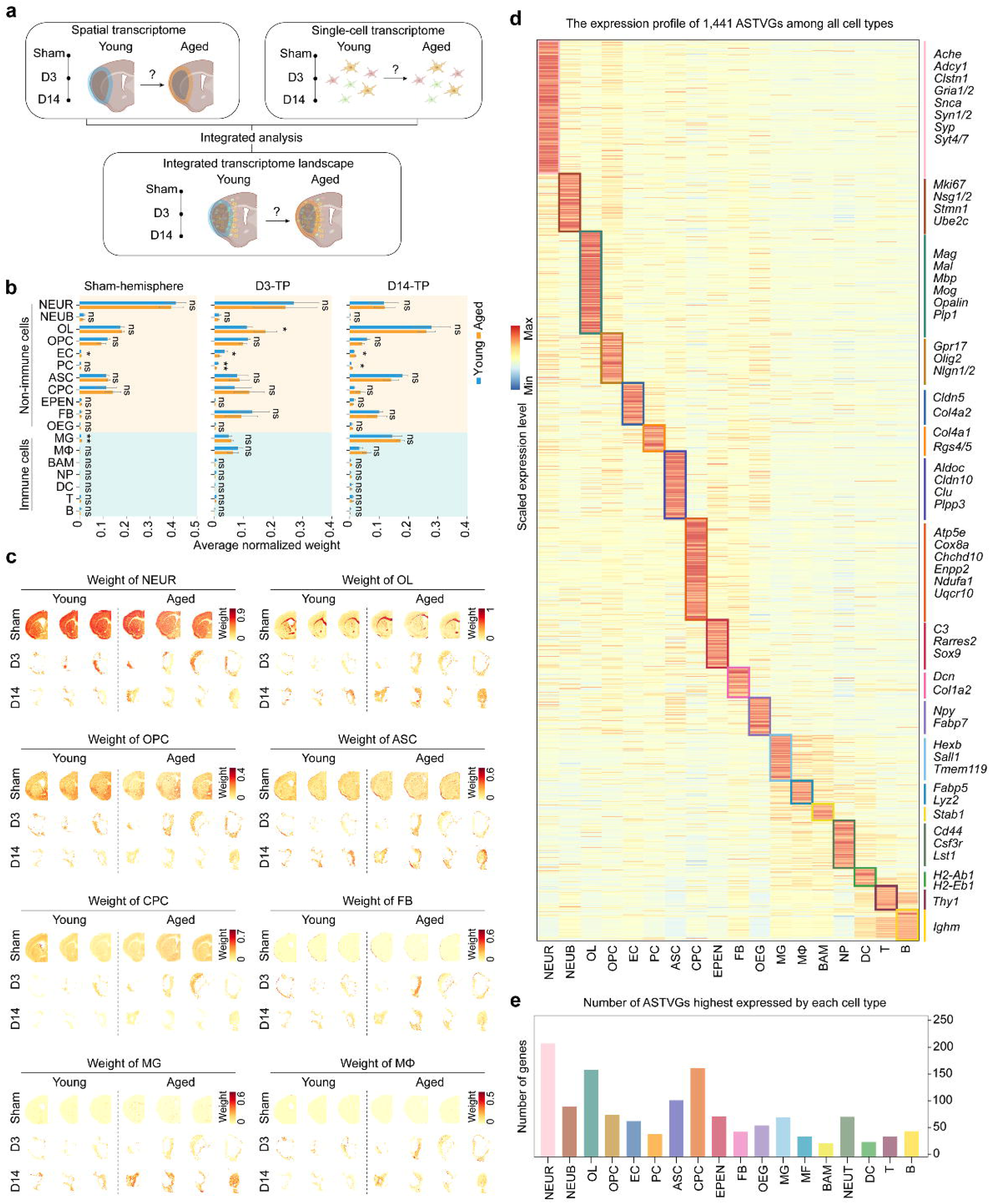
Age-dependent cellular profiling of age-sensitive time-varying genes by integrating spatial and single-cell transcriptomics in TP of ischemic stroke. **(a)** Schematic workflow for integrated analysis of spatial and single-cell transcriptome data. **(b)** Bar plots showing average normalized weights of 18 cell types in TP spots, quantified via RCTD deconvolution. **(c)** Spatial visualization of normalized RCTD weights for 8 dominant cell types enriched in TP regions. **(d)** Heatmap displaying scaled expression profiles of 1,441 ASTVGs across all 18 cell types in scRNA-seq data. **(e)** Bar plots showing the number of ASTVGs with cell-type-specific highest expression in each cell type. Cell-type weight quantification was performed using the RCTD deconvolution algorithm. ASTVGs: age-sensitive time-varying genes; ASC: astrocytes; B: B lymphocytes; BAM: border-associated macrophages; CPC: choroid plexus epithelial cells; DC: dendritic cells; D3: day 3 post-operation; D14: day 14 post-operation; EC: endothelial cells; EPEN: ependymal cells; FB: fibroblasts; MG: microglia; tMCAO: transient middle cerebral artery occlusion; MΦ: macrophages; NEUB: neuroblasts; NEUR: neurons; NP: neutrophils; OEG: olfactory ensheathing glia; OL: oligodendrocytes; OPC: oligodendrocyte precursor cells; PC: pericytes; RCTD: robust cell type decomposition; scRNA-seq: single-cell RNA sequencing; TP: transcriptomic penumbra; TVGs: time-varying genes.

Comparative analysis revealed minimal differences in TP cellular weights between young and aged (**Fig. 5b–c**), suggesting that age-dependent TP transcriptomic changes arise primarily from cell-intrinsic transcriptomic dysregulation rather than altered cell proportions. We further mapped the cellular source of 1,441 ASTVGs using scRNA-seq data (**Fig. 5d**), with NEUR, CPC, OL, ASC, and neuroblasts (NEUB) emerging as the top 5 enriched cell populations enriched for aging-sensitive genes (**Fig. 5e**).

Our previous scRNA-seq data contained a limited number of NEUR, likely due to neuronal loss during tissue dissociation, which precluded analysis of their age-dependent differences^24^. However, integrated spatial and single-cell data revealed the highest ASTVG enrichment in NEUR (∼200 ASTVGs), including core synaptic regulators (*Ache*, *Syp*, etc.) (**Fig. 5e**), which directly reflects the inherent age sensitivity of NEUR. OL also showed robust aging sensitivity (∼150 ASTVGs), including major myelin-encoding genes (*Mag*, *Mal*, etc.). CPC exhibited unexpectedly high aging susceptibility, suggesting impaired cerebrospinal fluid homeostasis in the aged TP. Among immune compartments, MG and neutrophils (NP) carried substantial aging-sensitive signatures, consistent with the literature ^47,48^.

To validate cellular origins of TP aging perturbations, we compared aging-induced DEGs in TP with cell-type-specific aging-induced DEGs at acute and chronic stages after stroke (**Supplementary Fig. 7**), revealing overlapping signatures across OL, MG, NP, and CPC, etc., thereby underscoring the contributions of these cell types to age-related transcriptomic differences in the TP.

Collectively, aging reshapes transcriptomic profiles across multiple cell types in the TP. Cell-intrinsic molecular dysregulation—rather than altered cell proportions—is the primary driver of the susceptibility of the TP to aging.

### Integrated spatial and single-cell transcriptomic decoding of aging-impaired pro-repair ligand-receptor networks in TP

Visium spots contain 1–10 cells, supporting ligand-receptor interactions within or between adjacent spots due to spatial proximity^49^ (**Fig. 6a**). CellChat analysis inferred 53 ligand-receptor pairs in TP, including age-specific, stage-specific, and conserved pairs (**Fig. 6b**). Given that loss of reparative ligand-receptor interactions contributes to age-related TP dysfunction, we focused on pairs unique to young TP but absent in aged TP. Acute-stage losses included TNR/FN1/COL4A2-SDC4, PTN-NCL, MAG-MAG, and GAS6-AXL pairs; chronic-stage losses included NRXN1/2/3-NLGN3, NEGR1-NEGR1, JAM3-JAM/F11R, FGF1-FGFR1/2/3, and CADM3-CADM3 pairs.

**Figure 6.**
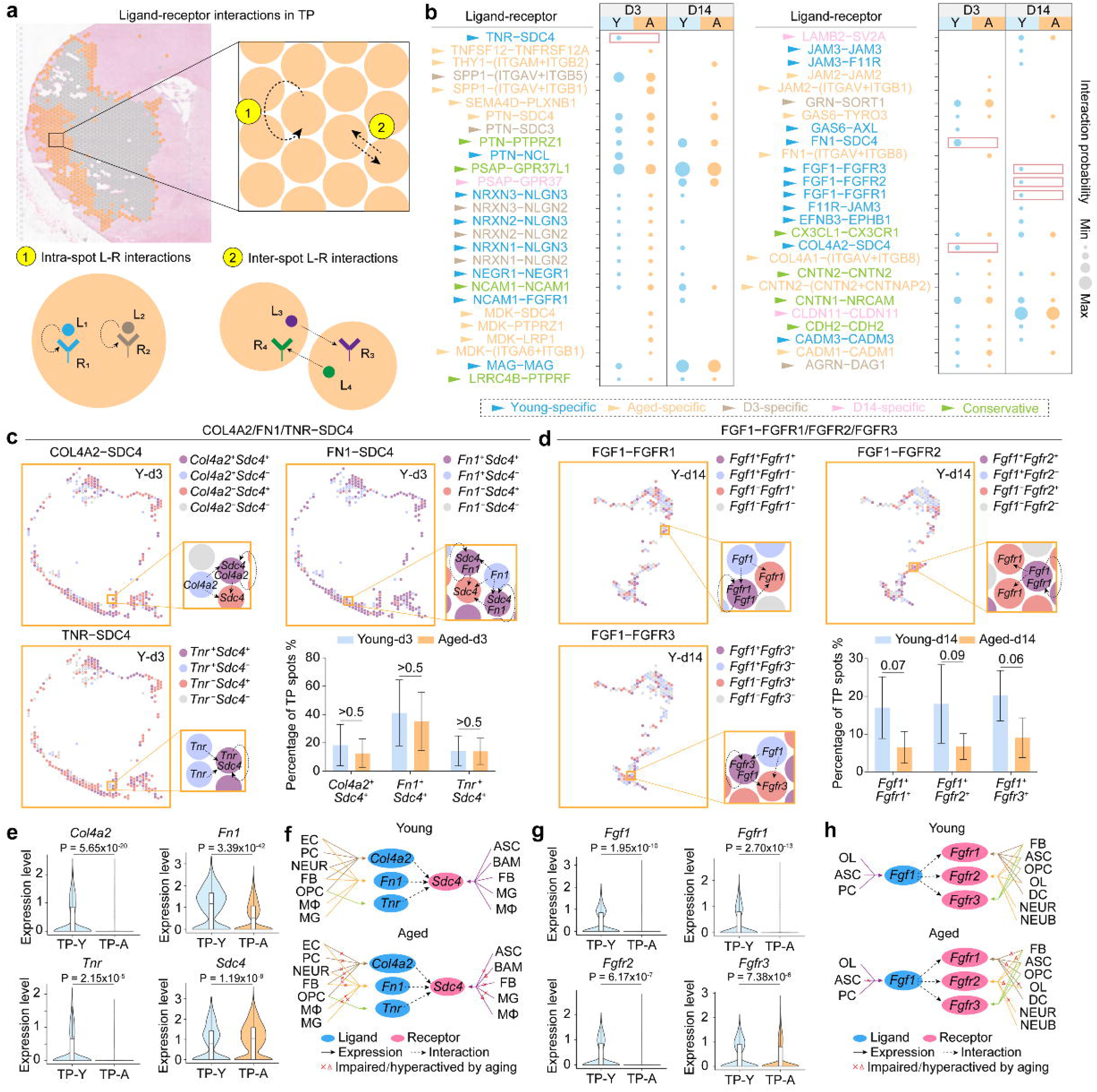
Age-dependent ligand-receptor interaction decoding by integrating spatial and single-cell transcriptomics in TP of ischemic stroke. **(a)** Schematic diagram illustrating the inference of intra-spot and inter-spot L-R interactions in TP regions. **(b)** Dot plots showing interaction probabilities of L-R pairs in TP across D3/D14 stages and young/aged groups. **(c–d)** Spatial visualization of spots expressing L-R pairs. Bar plots display the proportion of TP spots with L-R co-expression. **(e, g)** Violin plots showing expression differences of ligands and receptors between young and aged TP. Analysis of *TNR/FN1/COL4A2-SDC4* pairs is in (e); analysis of *FGF1-FGFR1/2/3* pairs is in (g). **(f, h)** Schematic diagram of cell-type-specific L-R pairing patterns and aging-related interaction impairment, inferred by integrating spatial and single-cell transcriptome data. . Analysis of *TNR/FN1/COL4A2-SDC4* pairs is in (f); analysis of *FGF1-FGFR1/2/3* pairs is in (h). L-R interaction prediction was conducted using the CellChat algorithm. *P*-values for inter-group expression comparisons were calculated by Wilcoxon rank-sum tests. ASC: astrocytes; BAM: brain-associated macrophages; D3: day 3 post-operation; D14: day 14 post-operation; EC: endothelial cells; EPEN: ependymal cells; FB: fibroblasts; L-R: ligand-receptor; MG: microglia; tMCAO: transient middle cerebral artery occlusion; MΦ: macrophages; OEG: olfactory ensheathing glia; OL: oligodendrocytes; OPC: oligodendrocyte precursor cells; TP: transcriptomic penumbra.

We focused on two core aging-impaired repair pathways, including acute-stage SDC4-related and chronic-stage FGF1-related pairs, both of which are established pro-repair genes^50,51^. Spatial co-expression analysis confirmed the simultaneous ligand-receptor co-expression in TP spots and strong spatial adjacency between ligand- and receptor-positive spots (**Fig. 6c–d**), robustly supporting the potential for intra- and inter-spot communication.

To clarify mechanisms underlying age-impaired interactions, we further analyzed age-dependent expression changes of core pathway genes. At the acute stage, *Col4a2*, *Fn1*, and *Tnr* were significantly downregulated in aged TP, whereas *Sdc4* expression was elevated, with no age-related change in ligand-*Sdc4* double-positive spot proportion (**Fig. 6c, e**). At the chronic stage, *Fgf1* and its receptors (*Fgfr1*–*3*) were markedly downregulated in the aged TP, with reduced *Fgf1*-receptor double-positive spot proportions (**Fig. 6d, g**). These findings suggest that acute *TNR*/*FN1*/*COL4A2*-*SDC4* impairment arises from ligand downregulation, whereas chronic *FGF1*-*FGFR1/2/3* dysfunction results from combined ligand-receptor downregulation and reduced co-expression. We also used the scRNA-seq data to map cellular sources of these ligands and receptors across aging (**Fig. 6f, h**). We further validated such age-dependent expression differences of several pro-repair ligand–receptor pairs at the protein level by immunofluorescence staining (**Supplementary Fig. 8**).

In summary, aging reshapes stage-specific TP molecular communication patterns, specifically abolishing multiple pro-repair ligand-receptor interactions via reduced ligand-receptor expression and co-expression.

### Integrated spatial and single-cell transcriptomic deciphering of core transcription factors for aging-driven dysfunction in the TP

Transcription factors (TFs) govern transcriptional programs underlying aging-related processes^52^. Accordingly, we performed single-cell regulatory network inference (SCENIC) analysis^53^ on scRNA-seq datasets from 18 cell types of young and aged mice at acute and chronic stages after stroke, establishing gene co-expression modules and gene regulatory networks (GRNs). TF activity was quantified via regulon activity (RA) scores (**Fig. 7a**). We screened age-sensitive TFs across all cell types by comparing RA between groups (|RA Young–RA Aged| > 0.1), obtaining cell-type-specific, stage-dependent, and age-sensitive TF landscapes (**Fig. 7b**, **Supplementary Table 13**).

**Figure 7.**
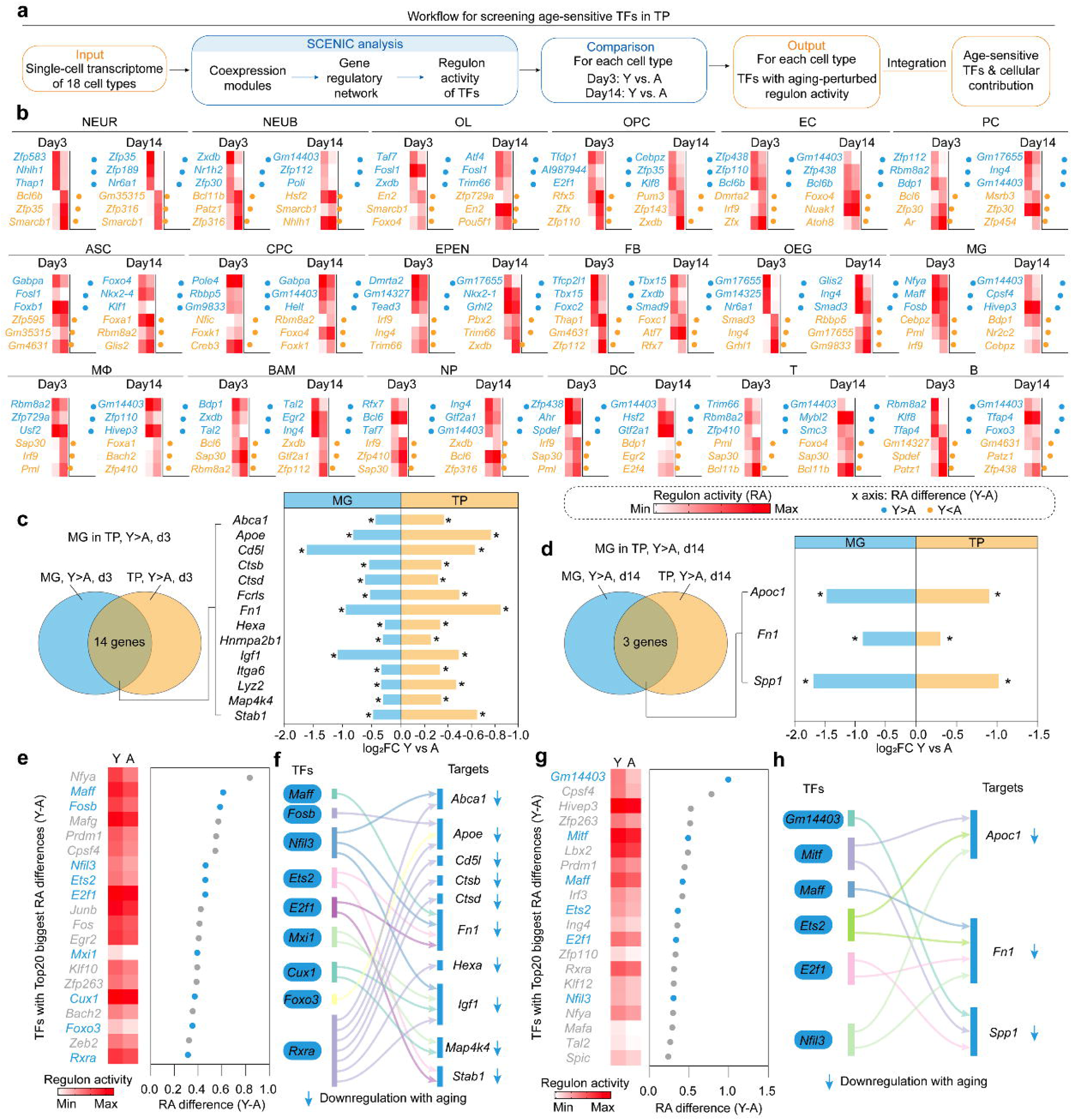
Age-dependent transcription factor regulon network deciphering by integrating spatial and single-cell transcriptomics in TP of ischemic stroke. **(a)** Schematic workflow illustrating the screening pipeline for age-sensitive TFs in TP via integrated spatial and single-cell transcriptomic analysis. **(b)** Heatmaps showing RA of representative age-sensitive TFs (top 3 positive & top 3 negative Y-A differences) across all cell types at D3 and D14 post-stroke. **(c– d)** Venn diagrams showing overlaps between aging-impaired genes in MG from scRNA-seq and TP aging-impaired genes from spatial transcriptomic data at acute D3 (c) and chronic D14 (d) stages. Bar plot showing the upregulaion level of these overlapped aging-impaired genes. *P*-values were calculated by Wilcoxon rank-sum tests with Bonferroni correction. **(e, g)** Ranking of top age-impaired TFs in MG at acute D3 (e) and chronic D14 (g) stages. Blue texts indicate overlapping upstream TFs of aging-impaired genes in (c, d). **(f, h)** Regulatory network diagrams showing TF-target gene regulatory relationships at acute D3 (f) and chronic D14 (h) stages. Regulon activity quantification was performed using the SCENIC algorithm. D3: day 3 post-operation; D14: day 14 post-operation; tMCAO: transient middle cerebral artery occlusion; MG: microglia; RA: regulon activity; scRNA-seq: single-cell RNA sequencing; TF: transcription factor; TP: transcriptomic penumbra.

Consistent with prior RCTD analysis and our previous findings^24^, MG represented major TP immune populations with age-related impairments in pro-repair functions. We therefore integrated spatial transcriptomic and scRNA-seq data to identify core upstream TFs driving aging-dependent MG dysfunction in TP. We overlapped aging-downregulated genes from MG scRNA-seq profiles with those from spatial transcriptomic profiles, identified 14 acute-stage aging-impaired genes (*Apoe*, *Cd5l*, etc.) and 3 chronic-stage aging-impaired genes (*Apoc1*, *Fn1,* and *Spp1*) (**Fig. 7c–d**), including key reparative mediators *Fn1* (encoding fibronectin)^54^, *Igf1* (encoding insulin-like growth factor-1)^55^, and *Spp1* (encoding osteopontin)^56^.

Upstream regulatory TFs for these overlapping target genes were predicted using the constructed GRNs (**Supplementary Table 14**). We further filtered these candidates against age-sensitive TFs in MG, ultimately pinpointing core TF drivers of age-dependent TP dysfunction (**Fig. 7e–h**). This analysis identified 9 acute-stage core TFs (*Maff, Fosb, Nfil3, Ets2, E2f1, Mxi1, Cux1, Foxo3*, and *Rxra*) (**Fig. 7f**) and 6 chronic-stage core TFs (*Gm14403, Mitf, Maff, Ets2, E2f1*, and *Nfil3*) (**Fig. 7h**).

In summary, using MG as a model cell type, we established an integrated pipeline for spatial transcriptomics and scRNA-seq to identify core TFs that drive cellular aging and functional deterioration in post-stroke TP.

## Discussion

We have established the first transcriptomic atlas of ischemic stroke as a function of age and time post-injury, comprising 20 young and aged mouse brains across sham, acute, and chronic post-stroke stages, with *in situ* profiling of 62,958 high-quality Visium spots. By integrating our new spatiotemporal transcriptome with matched scRNA-seq datasets, we defined and systematically characterized the TP, a spatially- and temporally resolved molecular entity in the ischemic peri-infarct region. Our work elucidates the dynamics of the TP across space and time, to reveal the molecular processes driven by aging and identify their associations with impaired penumbral repair, providing a comprehensive molecular foundation for therapeutics for ischemic stroke in the elderly.

Advances in single-cell and spatial transcriptomics have generated multiple datasets characterizing cellular responses after experimental ischemic stroke. Integrated spatial and single-cell transcriptomics have been used to dissect glial heterogeneity in young male mice with cerebral ischemia^25–27^. Another study conducted spatial transcriptomics on 19,777 spots coupled with scRNA in young photothrombotic stroke mice, revealing *Lgals9* (encodes Galectin-9) as a potential therapeutic target^23^. Nevertheless, these spatial transcriptomic studies focused solely on young animals, neglecting aging-related transcriptional changes. At the single-cell level, scRNA-seq profiling of dissociated tissues from young and aged tMCAO mice at 2 and 14 days post-stroke previously identified immune cell differences across age, but the mortality rate for aged males reached ∼90 % within three days post-injury (see Figure S14 in Garcia-Bonilla *et al.*^57^). Our aged males all survived until euthanasia at 14 days, permitting a full-factorial experimental design for the stroke mice. Our previous work used longitudinal scRNA-seq in young and aged tMCAO mice at 3 and 14 days, demonstrating that young mice initiate pro-repair transcriptional reprogramming in multiple cell types, whereas this endogenous compensatory effort is absent in aged mice^24^. In addition, reparative crosstalk between vascular and immune cells is severely impaired in aged brains, causing defective repair^24^. However, these studies lack the spatial resolution to specifically dissect molecular alterations within anatomical confines of the penumbra. Here, we fill this gap by constructing the first spatiotemporal landscape of the ischemic penumbra in young and aged mice across acute and chronic stages, integrating 62,958 high-quality spatial spots with matched scRNA-seq datasets. This dataset enables precise delineation of aging-driven TP alterations, offering a valuable public resource for elucidating mechanisms underlying age-dependent ischemic stroke outcomes.

The ischemic penumbra represents the salvageable peri-infarct tissue and core therapeutic target for clinical thrombolysis and thrombectomy, and the cornerstone of precision stroke care that enables extended therapeutic windows and targeted patient selection^8,11,12^. The penumbra was originally conceptualized by Astrup et al. in the 1970s, based on cerebral blood flow and electrophysiological thresholds^15,16^. Hossmann further refined this definition from a metabolic perspective^58^. Baron and Heiss then used PET imaging to establish the penumbra paradigm^59,60^, which was translated into the perfusion-diffusion MRI mismatch model by Schlaug and Warach^61,62^. Harston et al. improved penumbral imaging with pH-weighted MRI, showing that intracellular acidosis distinguishes the salvageable penumbra^63^. This physiological framework was extended to the molecular level by Sharp et al. via canonical molecular markers such as HSP70 ^18,64^. The neurovascular unit (NVU) concept redefined penumbral injury as a region of multicellular dysfunction involving neurons, glia, and vasculature^65^. Building on this, Lo et al. proposed a new penumbra model as a dynamic transition zone between injury and repair^17^. More recently, a multimodal mechanistic paradigm linked mitochondrial failure, neuroinflammation, and cell death to explain persistent translational failures in neuroprotection^66^. Thus, over the past four decades, penumbral research shifted from neurocentric views on protection to targeting the multicellular NVU, revealing marked spatial heterogeneity that warranted exploration beyond the classic penumbral concept. Despite these advances, spatially resolved molecular profiling of penumbra remained limited, restricting our understanding of the intrinsic molecular heterogeneity of peri-infarct regions. To fill this gap, we have defined the TP by integrating spatial transcriptomics with matched scRNA-seq datasets to systematically delineate the spatial molecular landscape of the penumbra and uncover its inherent periphery-to-core organization. We captured distinct stage-dependent biological shifts following ischemic injury. The acute stage was dominated by the stress response, metabolic disturbance, and excessive inflammatory activation, while the chronic phase was associated with progressive tissue remodeling, endogenous repair activation, and rewiring of intercellular communication. We further resolved stage-specific functional evolution, upstream transcriptional regulatory networks, and intercellular neurovascular-immune crosstalk across ischemic phases. This new framework advances our understanding of penumbral biology and provides promising molecular targets for therapeutic testing.

Aging remains the primary unchangeable risk factor determining pathological severity and long-term neurological prognosis after ischemic stroke. Compared with young individuals, elderly stroke patients exhibit more severe neurological impairment, reduced endogenous repair capacity, and limited responsiveness to clinical reperfusion interventions^67^, while the underlying molecular mechanisms remain incompletely elucidated. Based on our systematic analysis of the TP, aging exacerbates pathological deterioration during the acute ischemic phase, mainly characterized by disrupted energy homeostasis and impaired synaptic stability. These changes accelerate the irreversible progression of peri-infarct tissue. In the chronic repair phase, aging further suppresses endogenous reparative molecular programs, including those for white matter and vasculature in the TP, thereby triggering persistent immune activation and disrupting coordinated neurovascular-immune crosstalk^47^, which are essential for cerebral tissue remodeling^24,68^. Collectively, aging drives stage-dependent dynamic deterioration of the TP across acute and chronic stages. By weakening the intrinsic salvageable properties of the penumbra, aging directly worsens neurological outcomes in ischemic stroke. On the other hand, the current findings also identify potential compensatory changes in the aged TP, such as higher expression of markers of ATP synthesis/metabolism and aerobic respiration compared to the young TP on day 3 post-stroke (Figure 4e). At this early timepoint, there was also higher expression of neurogenesis, translation, and myelination markers in the aged TP (Figure 4e). Thus, it is tempting to speculate that aged mice would have suffered *even worse* neurological outcomes if these early gene expression changes in the TP were impeded with pharmacologic or genetic tools, thereby opening avenues for future mechanistic studies in aging animals.

In the present study, an increase in TP size was noted in aged mice by day 14 post-stroke (Figure S6b), but without considerable changes in cellular composition (Figure 5b). These unexpected findings based on gene expression can be contrasted with imaging data, in which the overall infarct size did not shift with aging. These discrepant imaging *versus* gene expression data reveal a need for spatial transcriptomic work in humans and suggest that RNA changes exert downstream effects that require additional time to unfold at the protein and structural/functional levels. Based on clinical and preclinical imaging data alone, the salvageable penumbra is believed to be smaller in size initially and disappear faster in the old, with a rapid transition into a dead infarct core. In juxtaposition to this widely held view, our spatial transcriptomic data suggest the continued persistence and even an expansion of a salvageable structure in the aged, indicating that the penumbra must be defined by multiple modalities—including early-stage mRNA changes—to reduce the knowledge barriers to treating the elderly in the future. Clinical data support this hopeful view, because even the elderly are known to benefit from thrombolytic treatment within the qualifying time window^69^. Given the expansion of the aged TP in the chronic injury stage, modulating expression of the gene targets identified herein could extend the time window for stroke treatment in the future. Hence, the present report opens multiple avenues for future exploration to further advance the field of aging-related stroke.

While our transcriptomic dataset provides a robust foundation, future studies will need to integrate proteomic and post-translational modification analyses to validate the correlation between transcriptomic alterations and functional phenotypes in pathological contexts. Second, the age- and stage-regulated pathways and candidate targets identified by our bioinformatic analysis offer clear directions for subsequent functional validation in cell culture and animal models, to confirm their biological roles and support clinical translation. Third, building on transcriptional reprogramming in the ischemic penumbra, future research can explore dynamic changes in the infarct core and non-injured tissue during stroke progression, alongside their impact on stroke pathogenesis and repair, as well as age-dependent regulatory mechanisms governing these regions. In a broader context, these age-specific molecular signatures and newly actionable candidates form a rational basis for targeted combination therapies. Thus, we propose that multiple molecular targets be manipulated together, for maximization of penumbral salvage, mitigation of aging-exacerbated brain injury, and improvement of long-term functional outcomes in elderly stroke patients.

In conclusion, we have capitalized on the original TP concept to systematically dissect molecular networks underlying age-dependent dysregulation of the ischemic penumbra. This study thus establishes the first publicly available spatiotemporal atlas of ischemic stroke spanning acute and chronic injury stages as a function of animal age, neural cell type, CNS region, and core-versus-penumbra relative location. These new datasets provide the field with a practical resource for subsequent preclinical and clinical testing of therapies that may salvage the dynamic penumbra in both young and old.

## Method

### 1. Animal

Young male C57BL/6J mice were purchased from The Jackson Laboratory and acclimatized for 2 weeks before experimentation, starting at 10–12 weeks of age. Aged male C57BL/6 mice were provided by the National Institute of Aging 1–2 months prior to experiments and acclimatized until use at 19–20 months of age. All mice were housed in a temperature- and humidity-controlled facility under a 12 h light/12 h dark cycle, with free access to standard chow and water ad libitum. All animal procedures were approved by the Institutional Animal Care and Use Committee of the University of Pittsburgh and conducted in accordance with the Guide for the Care and Use of Laboratory Animals. All measures were taken to minimize animal suffering and reduce the total number of animals used.

### 2. Experimental Stroke Model

Transient focal cerebral ischemia was induced by 60 min intraluminal occlusion of the left middle cerebral artery (MCA) as previously described^70^. Sham-operated mice received identical anesthesia and surgical exposure without MCA occlusion. Briefly, mice were initially anesthetized with 3% isoflurane in a gas mixture of 30% O / 67% N O until loss of tail-pinch reflex. Anesthesia was maintained throughout surgery with 1.5% isoflurane in 30% O / 68.5% N O via a nose cone under spontaneous breathing. A 7-0 silicon-coated monofilament was inserted into the common carotid artery, advanced to the MCA origin, and kept in place for 60 min to induce focal ischemia.

Rectal temperature was maintained at 37.0 ± 0.5 °C using a feedback-regulated heating pad during surgery. Regional cerebral blood flow (rCBF) was monitored by laser Doppler flowmetry (LDF) or two-dimensional laser speckle imaging. Mice exhibiting <70% reduction in rCBF during MCA occlusion relative to baseline were excluded from subsequent analyses. All surgical procedures and outcome quantifications were performed by investigators blinded to experimental group allocation.

### 3. MRI Assessment

T2-weighted MRI acquisition and quantitative analysis were performed by a blinded investigator. Mice were anesthetized with 1%–2% isoflurane delivered in an air/O mixture (2:1) via a nose cone. MRI scans were acquired on a 9.4 T/30 cm AVIII HD spectrometer (Bruker Biospin) equipped with a 12 cm high-performance gradient system, an 86 mm quadrature RF volume transmit coil, and a 2-channel surface receive coil, and operated with Paravision 6.0.1. A T2-weighted RARE sequence was applied with the following parameters: TR/TE = 4000/40 ms, FOV = 20 × 20 mm, acquisition matrix = 256 × 256, 21 contiguous slices at 0.5 mm thickness, 4 averages, RARE factor=8.

### 4. 10x Visium Spatial Transcriptome Profiling

#### 4.1 Tissue preparation, permeabilization and in situ capture

Fresh brain tissues were embedded in OCT compound and stored at −80 °C. Cryosections were cut at 10 μm thickness using a cryostat (Epredia HM525 NX) and mounted onto capture areas of Visium spatial gene expression slides. All tissue processing followed the 10x Genomics Visium Spatial Gene Expression protocol (CG000239 Rev F). Briefly, sections were fixed in methanol and stained with hematoxylin and eosin (H&E). Histological images were acquired under a Nikon Eclipse 90i microscope with a 10× objective. After imaging, tissue permeabilization was performed for 18 min to enable in situ mRNA capture onto spatial barcodes. Subsequent procedures, including spatially indexed cDNA reverse transcription and library construction, were conducted strictly following the manufacturer’s protocol.

#### 4.2 Library preparation and sequencing

Captured cDNA was eluted, denatured, and subjected to reverse transcription and second-strand synthesis. Library size distribution and concentration were evaluated using an Agilent 4150 TapeStation system. Libraries were sequenced on an Illumina platform with paired-end settings recommended by 10x Genomics, aiming for a sequencing depth of ≥50,000 read pairs per spot.

#### 4.3 Data preprocessing

Raw sequencing reads were processed using Space Ranger (v2.0.1, 10x Genomics). The workflow included sample demultiplexing, spatial barcode and UMI extraction from R1 reads, and alignment of R2 reads to the mm10 reference genome to generate spot-level gene expression matrices. Tissue images were co-registered with spatial spot coordinates for spatial mapping. Downstream analyses, encompassing expression normalization, dimensionality reduction, clustering and detection of spatially variable genes, were performed using Seurat in R and Scanpy in Python.

### 5. Transcriptome Data Analysis

#### 5.1 Basic processing of Visium spatial transcriptome data

Raw Visium data were processed using the Seurat package (v4.3.0)^71^ in R (v4.3.1). After dataset import, spot-level expression matrices were normalized and log-transformed. The top 5,000 highly variable genes were identified using FindVariableFeatures. Data were scaled with ScaleData, followed by principal component analysis for dimensional reduction. Unsupervised clustering was implemented using FindClusters. UMAP was applied for nonlinear dimensionality reduction and visualization via SpatialDimPlot. Spatial gene expression patterns were visualized using SpatialFeaturePlot and plotSurface from the SPATA2 package (v0.1.0)^72^.

#### 5.2 Basic processing of scRNA-seq data

scRNA-seq data were processed using Seurat (v4.3.0)^71^ in R (v4.3.1). Low-quality cells were filtered according to the following criteria: total UMI count <1,500 or >30,000; detected gene number <250 or >6,000; mitochondrial gene proportion >10%. Cells passing quality control were normalized, log-transformed, and scaled. The top 5,000 highly variable genes were selected, followed by PCA and unsupervised clustering using standard Seurat workflows.

#### 5.3 Differential expression analysis

Differentially expressed genes (DEGs) were identified using the Wilcoxon rank-sum test implemented in FindMarkers/FindAllMarkers. Genes with |log FC| > 0.25 and Bonferroni-adjusted P < 0.05 were defined as significant DEGs.

#### 5.4 Analysis of modularized temporal dynamics of TP

Temporal variable genes (TVGs) were defined as DEGs showing significant expression changes in at least one pairwise comparison among sham, day 3 post-stroke, and day 14 post-stroke TP. TVGs were partitioned into distinct modules based on their temporal expression patterns. For each module, the average expression trajectory across time points was calculated and plotted. Functional enrichment was performed for genes within each module; the top 50 significant GO terms were categorized into 6 major functional categories and ∼40 minor categories. For each functional subcategory and individual GO term, average gene expression profiles across time were generated and visualized. Within each term, the gene with the highest average expression was annotated as a representative signature gene, and its spatial expression was visualized in representative samples across time points.

#### 5.5 Analysis of aging-dependent perturbation on the temporal dynamics of TP

Age-sensitive TVGs (ASTVGs) were defined as TVGs exhibiting altered module assignment and temporal trajectories between young and aged mice. GO enrichment and functional classification were performed separately for ASTVGs in each module, with enriched terms grouped into 6 major categories and ∼40 minor categories. Temporal expression trajectories of functional subcategories and individual GO terms were compared between young and aged groups. Wilcoxon rank-sum tests were performed on spot-averaged gene expression at each time point to evaluate age-related differences. Spatial expression patterns of representative terms were further compared and visualized in matched young and aged samples at corresponding time points.

#### 5.6 Functional enrichment analysis

GO biological process enrichment analysis was performed using the Metascape online tool^73^. The entire mouse genome was used as the background gene set. Enrichment parameters were set to a minimum gene count of 3 and an enrichment factor > 1.5. Terms with P < 0.01 were considered statistically significant.

#### 5.7 Ligand-receptor interaction analysis

Cell–cell ligand–receptor communication across spatial spots was inferred using the CellChat R package (v1.6.1)^74,75^. Normalized expression matrices were imported, and overexpressed ligands/receptors were identified using default pipelines. Communication probability and pathway-level interaction networks were calculated with computeCommunProb and aggregateNet. Predicted ligand–receptor pairs and interaction strengths were visualized using ggplot2 (v3.4.4).

#### 5.8 Gene regulatory network inference by pySCENIC

Gene regulatory networks and regulon activity were inferred using pySCENIC (v0.12.1)^76^ on processed scRNA-seq matrices. The standard three-step workflow was applied: (1) TF–target co-expression modules were constructed using GRNBoost2; (2) cis-regulatory motif analysis was performed with cisTarget to filter indirect interactions and define high-confidence regulons; (3) regulon activity per cell was quantified using AUCell to generate AUC activity scores. Regulon activity matrices were exported as loom files for subsequent integration.

#### 5.9 Integration analysis with scRNA-seq: RCTD

Spatial cell-type deconvolution was performed using Robust Cell Type Deconvolution (RCTD)^45^ implemented in the spacexr R package (v2.2.1). Preprocessed scRNA-seq reference data, cell-type annotations, and UMI counts were compiled into RDS objects. Spatial transcriptomics data, including spot coordinates, expression matrices, and per-spot UMI counts, were similarly formatted. RCTD objects were constructed for each slice with max_cores=24, test_mode=F, and CELL_MIN_INSTANCE=6. Deconvolved cell-type weight matrices were normalized to ensure each spot summed to 1, and cell-type proportion profiles were integrated into downstream spatial analyses.

### 6. Immunofluorescence Staining and Imaging

Coronal brain sections (25 μm) were used for floating immunofluorescence staining. Sections were blocked with 5% donkey serum in 0.3% Triton X-100 PBS (PBST) for 1 h at room temperature, then incubated overnight with primary antibodies at 4 °C. After three washes in 0.3% PBST, sections were incubated with fluorophore-conjugated secondary antibodies for 1 h at room temperature. For double immunostaining, the incubation and washing steps were repeated sequentially. Sections were mounted with DAPI-containing Fluoromount-G. For mouse-derived primary antibodies, the M.O.M. Immunodetection Kit (Vector Laboratories) was used to reduce nonspecific background according to the manufacturer’s protocol. Primary antibody information is provided (**Supplementary Table 15)**. All secondary antibodies were diluted to 1:1000.

Whole-section imaging was acquired using the EVOS M7000 system, and high-resolution confocal images were captured with a Nikon A1 microscope. For quantitative analysis, one to two random fields in the peri-infarct region were selected per section; two anatomical sections covering the infarct territory were analyzed per mouse. Image quantification was performed in ImageJ by two independent investigators blinded to group allocation. Positively stained cells were manually annotated to avoid duplicate counting. The infarct core was defined by condensed DAPI nuclear morphology. The peri-infarct penumbra region was defined as the tissue extending 200–300 μm radially from the infarct border.

## Data Availability

All transcriptomic data in this study can be download and interactively accessed via Single Cell Portal at https://singlecell.broadinstitute.org/ (SCP3686 for scRNA-seq and SCP3687 for Visium spatial transcriptome. Other data generated during the study are available from the authors upon reasonable request.

## Acknowledgement

This work was supported by the following grants: AHA Transformational Award (969858), the VA SRCS Award (821-RC-NB-30556), VA Merit Review grants I01 BX005290 and I01 BX003377 (to J.C.). J.C. is also supported, in part, by Richard K. Mellon Endowed Chair from Department of Neurology at the University of Pittsburgh School of Medicine. H.P. is supported in part by the AHA Career Development Award (926806).

## Author contributions

J.C. conceived, designed, and supervised the study. C.J., L.F., Q.Y., H.P., W.Z., J.X. L., and W-C.W. performed experiments and data collection. C.J. and G.H. conducted data analysis. J.C. and C.J. performed data interpretation and literature review. C.J., Y.S., and K.C. performed resource data processing and management. C.J. prepared the figures on resource data; H.P., W.Z. and J.X. L. prepared the figures on immunofluorescence. C.J. and J.C. drafted the manuscript. Y.S., R.K.L., X.H., and K-J. Y. revised the manuscript. All authors contributed to the review of the manuscript and approved the final version.

## Supplementary information

**Supplementary Figure 1. T2-weighted MRI validation of anatomical location and infarct lesion in all enrolled samples.** Grayscale coronal T2-weighted MRI images of all ischemic mice at day 3 after tMCAO. The imaging plane matches the anatomical position of Visium tissue sections used for spatial transcriptomics. D3: day 3 post-operation; tMCAO: transient middle cerebral artery occlusion; MRI: magnetic resonance imaging.

**Supplementary Figure 2. Robust transcriptional distinction among spatial clusters and ribosomal pathway enrichment and gene expression in Les3 and Les14.** (a) Heatmap showing expression patterns of the top 10 marker genes (ranked by logFC) across seven spatial clusters (Ctx, Str, WM, PV, SN, Les3, Les14) in all spatial spots. (b) Bar plot showing ribosome-associated functional enrichment terms of upregulated genes in Les3. (c) Bar plot showing expression levels of representative ribosomal genes enriched in Les3. (d) Bar plot showing ribosome-associated functional enrichment terms of upregulated genes in Les14. (e) Bar plot showing expression levels of representative ribosomal genes enriched in Les14. Les3: lesion region at day 3 post-stroke; Les14: lesion region at day 14 post-stroke; logFC: log2 fold-change.

**Supplementary Figure 3. Spatial localization and transcriptomic specificity of Peri subclusters surrounding the infarct core.** (a) UMAP plot showing five subclusters (Core1–3, Peri1–2) derived from re-clustering of Les3. (b) UMAP plot showing four subclusters (Core1–2, Peri1–2) derived from re-clustering of Les14. (c) SpatialPlot showing anatomical distribution of five Les3 subclusters across acute-stage samples. (d) SpatialPlot showing anatomical distribution of four Les14 subclusters across chronic-stage samples. (e) From left to right: infarct area defined by H&E staining, infarct area defined by T2- weighted MRI, spatial distribution of Peri spots, and multimodal aligned composite image. (f) Number of differentially expressed genes between Peri and Core subpopulations. Core: infarct core subcluster; H&E: hematoxylin and eosin; Les3: lesion region at day 3 post-stroke; Les14: lesion region at day 14 post-stroke; MRI: magnetic resonance imaging; Peri: penumbral subcluster; UMAP: uniform manifold approximation and projection.

**Supplementary Figure 4. Screening and spatial expression validation of candidate TP marker genes.** (a) Spatial expression of 30 acute-phase TP marker genes across all acute-stage samples. (b) Spatial expression of 70 chronic-phase TP marker genes across all chronic-stage samples. Green lines indicate raw expression levels; orange lines indicate smoothed expression processed by the SPATA2 package. TP: transcriptomic penumbra.

**Supplementary Figure 5. Protein-level validation of age-conserved TP marker expression across acute and chronic stroke stages.** (a) Representative immunofluorescence scans showing the expression of GFAP, DHRS1, and FABP7 in the acute TP (day 3 post-stroke) in young versus aged mice. (b) Representative immunofluorescence scans showing the expression of GFAP, FXYD1, and PRDX6 in the chronic TP (day 14 post-stroke) in young versus aged mice. Left: ipsilateral (ischemic) hemisphere; Right: contralateral hemisphere.

**Supplementary Figure 6. Age-dependent alterations in TP size.** (a) Brown mask showing the range of TP area and the range of ipsilesional hemisphere in each sample. (b) Bar plots showing the proportion of TP area or the density of TP spots in the ipsilesional hemisphere among groups. TP: transcriptomic penumbra.

**Supplementary Figure 7. Integrated spatial and single-cell transcriptomic dissection of cellular contributions to age-dependent transcriptional alterations in TP.** (a) Overlap of age-related DEGs derived from spatial transcriptomic TP data and cell-type-specific age-related DEGs from scRNA-seq at acute and chronic stages. (b) Statistical summary of overlapping age-related DEG numbers across cell types in the chronic stage. DEG: differentially expressed gene; scRNA-seq: single-cell RNA sequencing; TP: transcriptomic penumbra.

**Supplementary Figure 8. Protein-level validation of age-impaired pro-repair ligand-receptor pairs in the post-stroke brain.** (a) Representative immunofluorescence images showing FN1 and SDC4 expression in the peri-infarct region at 3 days post-stroke in young versus aged mice, with FN1 co- localized with the endothelial marker CD31. (b) Representative immunofluorescence images showing FGF1 and FGFR1 expression in the peri-infarct region at 14 days post-stroke in young versus aged mice, with FGF1 co-localized with the astrocytic marker GFAP.

**Supplementary Table 1.** Sample information of 20 Visium spatial transcriptomic datasets.

**Supplementary Table 2.** Marker genes and functional enrichment of seven main spatial clusters.

**Supplementary Table 3.** Candidate TP marker genes identified by DEG analysis of TP versus Core and other brain regions.

**Supplementary Table 4.** TVGs in TP of young mice.

**Supplementary Table 5.** Module assignment of temporal variable genes in young mouse TP.

**Supplementary Table 6.** Functional enrichment of temporal modules in young mouse TP.

**Supplementary Table 7.** Age-dependent DEGs in TP and corresponding functional enrichment results.

**Supplementary Table 8.** TVGs in TP of aged mice.

**Supplementary Table 9.** Module assignment of temporal variable genes in aged mouse TP.

**Supplementary Table 10.** Module transition characteristics of aging-sensitive time-varying genes (ASTVGs).

**Supplementary Table 11.** Functional enrichment of temporal modules of ASTVGs in aged vs young TP.

**Supplementary Table 12.** Cell composition weights of TP inferred by RCTD.

**Supplementary Table 13.** Transcription factor regulon activity across cell types at acute and chronic stages based on pySCENIC analysis.

**Supplementary Table 14.** Regulatory relationships between transcription factors and downstream target genes inferred by pySCENIC.

**Supplementary Table 15.** Information of primary antibodies for immunofluorescence.

